# Identification of a Pangolin Niche for a 2019-nCoV-like Coronavirus via an Extensive Meta-metagenomic Search

**DOI:** 10.1101/2020.02.08.939660

**Authors:** Lamia Wahba, Nimit Jain, Andrew Z. Fire, Massa J. Shoura, Karen L. Artiles, Matthew J. McCoy, Dae-Eun Jeong

**Affiliations:** Department of Pathology, Stanford University School of Medicine, Stanford, CA 94305, USA; Department of Genetics, Stanford University School of Medicine, Stanford, CA 94305, USA; Department of Bioengineering, Stanford University, Stanford, CA 94305, USA

**Author notes:** Co-corresponding and equally contributing authors. Author order was chosen randomly. **Author for publication correspondence:** Andrew Fire.

## Abstract

In numerous instances, tracking the biological significance of a nucleic acid sequence can be augmented through the identification of environmental niches in which the sequence of interest is present. Many metagenomic datasets are now available, with deep sequencing of samples from diverse biological niches. While any individual metagenomic dataset can be readily queried using web-based tools, meta-searches through all such datasets are less accessible. In this brief communication, we demonstrate such a meta-meta-genomic approach, examining close matches to the Wuhan coronavirus 2019-nCoV in all high-throughput sequencing datasets in the NCBI Sequence Read Archive accessible with the keyword “virome”. In addition to the homology to bat coronaviruses observed in descriptions of the 2019-nCoV sequence (F. Wu et al. 2020, Nature, doi.org/10.1038/s41586-020-2008-3; P. Zhou et al. 2020, Nature, doi.org/10.1038/s41586-020-2012-7), we note a strong homology to numerous sequence reads in a metavirome dataset generated from the lungs of deceased Pangolins reported by Liu et al. (Viruses 11:11, 2019, http://doi.org/10.3390/v11110979). Our observations are relevant to discussions of the derivation of 2019-nCoV and illustrate the utility and limitations of meta-metagenomic search tools in effective and rapid characterization of potentially significant nucleic acid sequences.

**Importance:** Meta-metagenomic searches allow for high-speed, low-cost identification of potentially significant biological niches for sequences of interest.

## Introduction

In the early years of nucleic acids sequencing, aggregation of the majority of published DNA and RNA sequences into public sequence databases greatly aided biological hypothesis generation and discovery. Search tools capable of interrogating the ever-expanding databases were facilitated by creative algorithm development and software engineering, and by the ever-increasing capabilities of computer hardware and the internet. As of the early 2000s, sequencing methodologies and computational technologies advanced in tandem, enabling quick homology results from a novel sequence without substantial cost.

With the development of larger-scale sequencing methodologies, the time and resources to search all extant sequence data became untenable for most studies. However, creative approaches involving curated databases and feature searches ensured that many key features of novel sequences remained readily accessible. At the same time, the nascent field of metagenomics began, with numerous studies highlighting the power of survey sequencing of DNA and RNA from samples as diverse as the human gut and Antarctic soil (1, 2). As the diversity and sizes of such datasets expand, the utility of searching them with a novel sequence increases. Meta-metagenomic searches are currently underutilized. In principle such searches would involve direct access to sequence data from a large set of metagenomic experiments on a terabyte scale, along with software able to search for similarity to a query sequence. We find that neither of these aspects of meta-metagenomic searches is infeasible with current data transfer and processing speeds. In this communication, we report the results of searching the recently-described 2019-nCoV coronavirus sequence through a set of metagenomic datasets with the tag “virome”.

## Materials and Methods

### Computing Hardware

A Linux workstation used for the bulk analysis of metagenomic datasets employs an 8-core i7 Intel Microprocessor, 128G of Random Access Memory, 12TB of conventional disk storage, and 1TB of SSD storage. Additional analyses of individual alignments were carried out with standard consumer-grade computers.

### Sequence data

All sequence data for this analysis were downloaded from the National Center for Biotechnology Information (NCBI) website, with individual sequences downloaded through a web interface and metagenomic datasets downloaded from the NCBI Sequence Read Archive (SRA) using the SRA-tools package (version 2.9.1). The latter sequence data were downloaded as .sra files using the prefetch tool, with extraction to readable format (.fasta.gz) using the NCBI fastq-dump tool. Each of these manipulations can fail some fraction of the time. Obtaining the sequences can fail due to network issues, while extraction in readable format occasionally fails for unknown reasons. Thus the workflow continually requests .sra files with ncbi-prefetch until at least some type of file is obtained, followed by attempts to unpack into .fasta.gz format until one such file is obtained from each .sra file. Metagenomic datasets for analysis were chosen through a keyword search of the SRA descriptions for “virome” and downloaded between January 27 and January 31, 2020. We note that the “virome” keyword search will certainly not capture every metagenomic dataset with viral sequences, and likewise not capture every virus in the short sequence read archive. With up to 16 threads running simultaneously, total download time (prefetch) was approximately 2 days. Similar time was required for conversion to gzipped fasta files. A total of 9014 sequence datasets were downloaded and converted to fasta.gz files. Most files contained large numbers of reads with a small fraction of files containing very little data (only a few reads or reads of at most a few base pairs). The total dataset consists of 2.5TB of compressed sequence data corresponding to approximately 10^13^ bases.

### Search Software

For rapid identification of close matches among large numbers of metagenomic reads, we used a simple dictionary based on the 2019-nCoV sequence (NCBI MN908947.3Wuhan-Hu-1) and its reverse complement, querying every 8th k-mer along the individual reads for matches to the sequence. As a reference, and to benchmark the workflow further, we included several additional sequences in the query (Vaccinia virus, an arbitrary segment of a flu isolate, the full sequence of bacteriophage P_4_, and a number of putative polinton sequences from *Caenorhabditis briggsae*). The relatively small group of k-mers being queried (<10^6^) allows a rapid search for homologs. This was implemented in a Python script run using the PyPy accelerated interpreter. We stress that this is by no means the most comprehensive or fastest search for large datasets. However, it is more than sufficient to rapidly find any closely matching sequence (with the downloading and conversion of the data, rather than the search, being rate limiting).

### Alignment of reads to 2019-nCoV

Reads from the positive pangolin datasets were adapter-trimmed with cutadapt (version 1.18) (3), and mapped to the 2019-nCoV genome with BWA-MEM (version 0.7.12) (4) using default settings for paired-end mode. Alignments were visualized with the Integrated Genomics Viewer IGV tool (version 2.4.10) (5).

### Assessment of nucleotide similarity between 2019-nCoV, pangolin metavirome reads, and closely related bat coronaviruses

All pangolin metavirome reads that aligned to the 2019-nCoV genome with BWA-MEM after adapter trimming with cutadapt were used for calculation. The bat coronavirus genomes were aligned to the 2019-nCoV genome in a multiple sequence alignment using the web-interface for Clustal Omega (https://www.ebi.ac.uk/Tools/msa/clustalo/) (6) with default settings. We note that sequence insertions with respect to the 2019-nCoV genome in either the pangolin metavirome reads or the bat coronavirus genomes are not accounted for in the similarity traces shown in Figure 1b.

**Figure 1.**
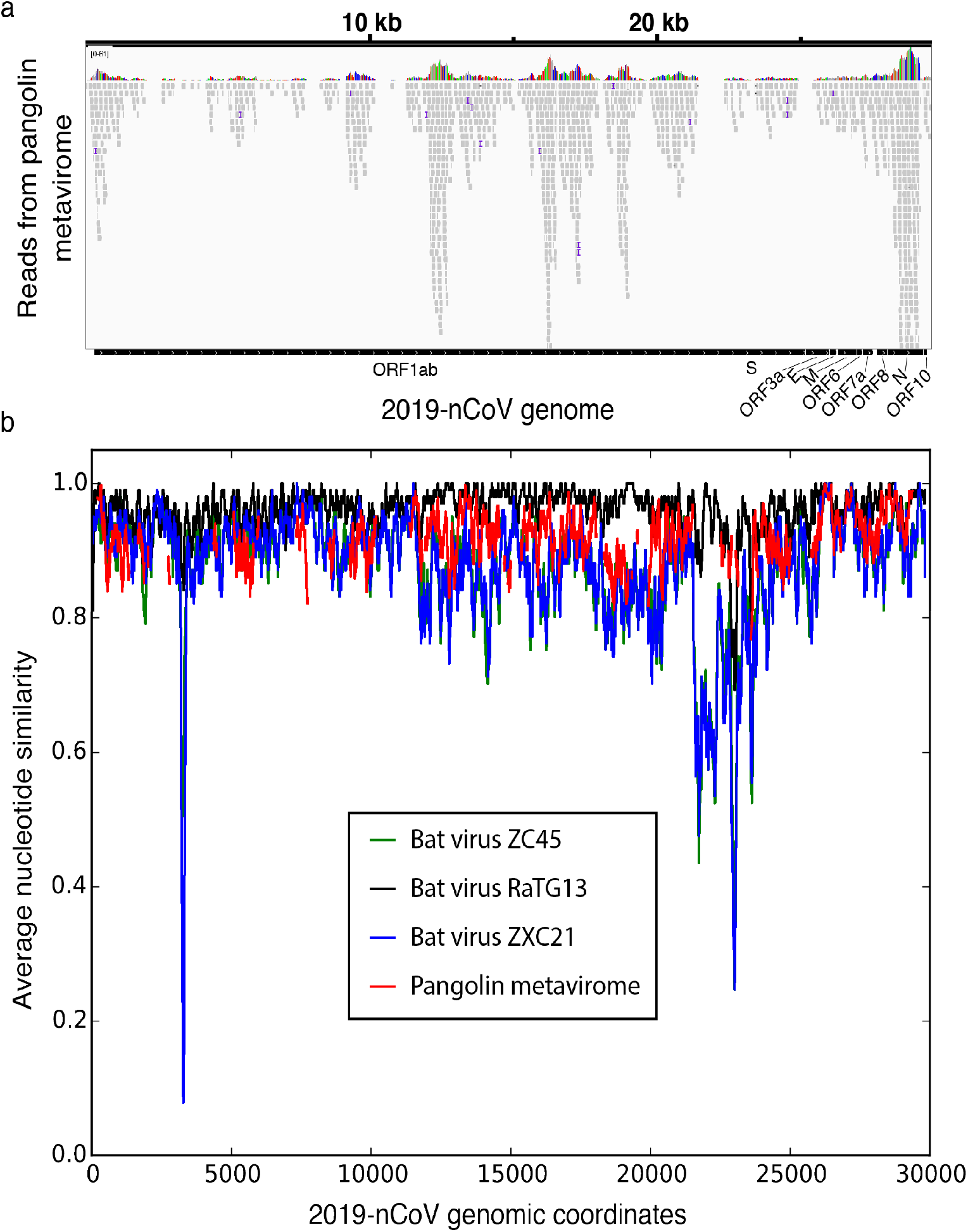
**(a) Integrated Genomics Viewer (IGV) snapshot of alignment.** Reads from the pangolin lung virome samples (SRR10168377, SRR10168378 and SRR10168376) were mapped to a 2019-nCoV reference sequence (Genbank accession: MN908947.3). The total number of aligned reads from the three samples was 1107, 313 and 32 reads respectively. **(b) Quantification of nucleotide-level similarity between the 2019-nCoV genome and pangolin lung metavirome reads aligning to the 2019-nCoV genome.** Average similarity was calculated in 101-nucleotide windows along the 2019-nCoV genome, and is only shown for those windows where each nucleotide in the window had coverage ≥ 2. Average nucleotide similarity calculated (in 101-nucleotide windows) between the 2019-nCoV genome and reference genomes of three relevant bat coronaviruses (bat-SL-CoVZC45: accession MG772933.1, bat-SL-CoVZXC21: accession MG772934.1, RaTG13: accession MN996532.1) is also shown. Note that the Pangolin metavirome similarity trace is not directly comparable to the bat coronavirus similarity traces, because the former uses read data for calculation whereas the latter use reference genomes.

### Regional assessment of synonymous and nonsynonymous mutations (Figure S1)

Although the incomplete nature of coverage in the Pangolin metavirome data somewhat limits the application of measures such as normalized dN/dS values, it remains possible to identify regions with the strongest matches of this inferred viral sequence with the human and bat homologs and to determine the distribution of synonymous and nonsynonymous variants in these regions. Details of this analysis are presented in Figure S1.

### Accessibility of Software

Scripts used for the observations described in this communication are available at https://github.com/firelabsoftware/Metasearch2020.

## Results

To identify biological niches that might harbor viruses closely related to 2019-nCoV, we searched through publicly available metaviromic datasets. We were most interested in viruses with highly similar sequences, as these would likely be most useful in forming hypotheses about the origin and pathology of the recent human virus. We thus set a threshold requiring matching of a perfect 32-nucleotide segment with a granularity of 8 nucleotides in the search (i.e., interrogating the complete database of k-mers from the virus with k-mers starting at nucleotide 1, 9, 17, 25, 33 of each read from the metagenomic data for a perfect match). This would catch any perfect match of 39 nucleotides or greater, with some homologies as short as 32 nucleotides captured depending on the precise phasing of the read.

All metagenomic datasets with the keyword “virome” in NCBI SRA as of January 2020 were selected for analysis in a process that required approximately 2 days each for downloading and conversion to readable file formats and one day for searching by k-mer match on a desktop workstation computer (i7 8-core). Together the datasets included information from 9014 NCBI Short Read Archive entries with (in total) 6.2*10^10^ individual reads and 8.4*10^12^ base pairs. Despite the relatively large mass of data, the 32-nucleotide k-mer match remains a stringent measure, with spurious matches to the ~30 kb 2019-nCoV genome expected at only 1 in 3*10^14^. Positive matches among the metagenomic datasets analyzed were relatively rare, with the vast majority of datasets (8994/9014 or 99.8%) showing no matched 32-mers to 2019-nCoV. Of the datasets with matched k-mers, one was from an apparent synthetic mixture of viral sequences, while the remaining were all from vertebrate animal sources. The matches were from five studies: two bat-focused studies (7, 8), one bird-focused study (9), one small-animal-and-rodent focused study (10), and a study of pangolins (11) [Table 1].

**Table 1.** Metagenomic datasets with k=32-mer matches to MN908947.3 (2019-nCoV). Details of search are described in legend to Table S1.

| Description | File | TotalReads | TotalBases | HitReads | HitKmers |
| --- | --- | --- | --- | --- | --- |
| Pangolin Lung | SRR10168376_1 | 18067615 | 2710142250 | 5 | 9 |
| Pangolin Lung | SRR10168376_2 | 18067615 | 2710142250 | 6 | 21 |
| Pangolin Lung | SRR10168377_1 | 16414925 | 2462238750 | 308 | 955 |
| Pangolin Lung | SRR10168377_2 | 16414925 | 2462238750 | 285 | 904 |
| Pangolin Lung | SRR10168378_1 | 19045923 | 2856888450 | 91 | 352 |
| Pangolin Lung | SRR10168378_2 | 19045923 | 2856888450 | 96 | 337 |
| Pangolin Lung | SRR10168392_1 | 39738679 | 5960801850 | 2 | 4 |
| Pangolin Lung | SRR10168392_2 | 39738679 | 5960801850 | 4 | 6 |
| "Mock virome" | SRR3458564_2 | 10957589 | 1654595939 | 1 | 1 |
| Bat Feces | SRR5040897_1 | 4020145 | 607041895 | 5 | 5 |
| Bat Feces | SRR5040897_2 | 4020145 | 607041895 | 4 | 4 |
| Bat Feces | SRR5040918_1 | 4804340 | 725455340 | 4 | 4 |
| Bat Feces | SRR5040918_2 | 4804340 | 725455340 | 3 | 3 |
| Virome analysis of Rodents at | SRR5343975_1 | 19775296 | 1582023680 | 1 | 1 |
| Virome analysis of Rodents at | SRR5343977_1 | 20227246 | 1618179680 | 1 | 2 |
| Bat | SRR5351751_2 | 363186 | 32644600 | 1 | 1 |
| Bat | SRR5351752_1 | 305206 | 27328640 | 24 | 57 |
| Bat | SRR5351752_2 | 305206 | 27608440 | 20 | 55 |
| Bat | SRR5351758_2 | 301889 | 27514580 | 1 | 3 |
| Bat | SRR5351760_1 | 788212 | 70634120 | 407 | 962 |
| Bat | SRR5351760_2 | 788212 | 71244040 | 378 | 1006 |
| Virome analysis of Rodents at | SRR5365807_1 | 21646498 | 1731719840 | 2 | 5 |
| Virome analysis of Rodents at | SRR5365809_1 | 22799703 | 1823976240 | 1 | 1 |
| Virome analysis of Rodents at | SRR5431767_1 | 10213803 | 1031594103 | 2 | 2 |
| Virome analysis of Rodents at | SRR5447167_1 | 13849913 | 1398841213 | 2 | 2 |
| Virome analysis of Rodents at | SRR5447174_1 | 14063346 | 1420397946 | 1 | 1 |
| Virome analysis of Rodents at | SRR5447175_1 | 7510256 | 758535856 | 1 | 1 |
| Avocet (RNA Virome in Wild f | SRR7239364_1 | 22336985 | 2233698500 | 2 | 2 |
| Metagenomic datasets with k=32-mer matches to MN908947.3 2019-nCoV<br>Details of search and are described in legend to Table S1 |  |  |  |  |  |

The abundance and homology of viruses within a metagenomic sample are of considerable interest in interpreting possible characteristics of infection and relevance to the query virus. From the quick k-mer search, an initial indicator could be inferred from the number of matching reads and k-mer match counts for those reads [Table 1, Supplementary Table 1]. For the 2019-nCoV matches amongst the available metagenomic datasets, the strongest and most abundant matches in these analyses came from the pangolin lung metaviromes. The matches were observed throughout the 2019-nCoV query sequence and many of the matching reads showed numerous matching 32-mer sequences. The vast majority of matches were in two lung samples with small numbers of matches in two additional lung datasets (11). No matches were detected for five additional lung datasets, and no matches were seen in eight spleen samples and a lymph node sample (11). Further analysis of coverage and homology through alignment of the entire metagenomic datasets revealed an extensive, if incomplete, coverage of the 2019-nCoV genome [Figure 1a]. Percent nucleotide similarity can be calculated for pangolin metavirome reads aligning to 2019-nCoV [Figure 1b], and these segmental homologies consistently showed strong matches, approaching (but still overall weaker than) the similarity of the closest known bat coronavirus (RaTG13).

## Discussion

Meta-meta-genomic searching can provide unique opportunities to understand the distribution of nucleic acid sequences in diverse environmental niches. As metagenomic datasets proliferate and as both the need and capability to identify pathogenic agents through sequencing increase, meta-metagenomic searching may prove extremely useful in tracing the origins and spreading of causative agents. In the example we present in this paper, such a search identifies a number of niches with sequences matching the genome of the recent 2019-nCoV virus. These analyses raise a number of relevant points for the origin of 2019-nCoV. Before describing the details of these points, however, it is important to stress that while environmental, clinical, and animal-based sequencing is valuable in understanding how viruses traverse the animal ecosphere, static sequence distributions cannot be used to construct a virus’ full transmission history amongst different biological niches. So even were the closest relative of a virus causing disease in species X to be found in species Y, we cannot define the source of the outbreak, or the direction(s) of transmission. As some viruses may move more than once between hosts, the sequence of a genome at any time may reflect a history of selection and drift in several different host species. This point is also accentuated in the microcosm of our searches for this work. When we originally obtained the 2019-nCoV sequence from the posted work of Wu et al., we recapitulated their result that bat-SL-CoVZC45 was the closest related sequence in NCBI’s non-redundant (nr/nt) database. In our screen of metavirome datasets, we observed several pangolin metavirome sequences—which are not currently in the (nr/nt) database—and which are more closely related to 2019-nCoV than bat-SL-CoVZC45. An assumption that the closest relative of a sequence identifies the origin would at that point have transferred the extant model to zoonosis from pangolin instead of bat. To complicate such a model, an additional study from Zhou et al. (12) described a previously unpublished Coronavirus sequence, designated RaTG13 with much stronger homology to 2019-nCoV than either bat-SL-CoVZC45 or the pangolin reads from Liu et al (11). While this observation certainly shifts the discussion (legitimately) toward a possible bat-borne intermediate in the chain leading to 2019-nCoV, it remains difficult to determine if any of these are true intermediates in the chain of infectivity.

The match of 2019-nCoV to the pangolin coronavirus sequences also enables a link to substantial context on the pangolin samples from Liu et al. (11), with information on the source of the rescued animals (from smuggling activity), the nature of their deaths despite rescue efforts, the potential presence of both other coronaviruses and other non-corona viruses in the same cells, and the accompanying pathology. That work describes analyses indicating several coronaviruses present in two of the pangolin lungs as well as other viral species in those lungs. The pangolins appear to have died from lung-related illness, which may have involved the 2019-nCoV closely-related virus. Notably, however, two of the deceased pangolin lungs had much lower coronavirus signals, while five showed no signal, with sequencing depths in the various lungs roughly comparable. Although it remains possible that the 2019-nCoV-like coronavirus was the primary cause of death for these animals, it is also possible (as noted by Liu et al. (11)) that the virus was simply present in the tissue, with mortality due to another virus, a combination of infectious agents, or other exposures.

During the course of this work, the homology between 2019-nCoV and pangolin coronavirus sequences in a particular genomic subregion was also noted and discussed in an online forum (“Virological.org”) with some extremely valuable analyses and insights.

Matthew Wong and colleagues bring up the homology to the pangolin metagenomic dataset in this thread and appear to have encountered it through a more targeted search than ours (this study has since been posted online in bioRxiv) (13). As noted by Wong et al. (13), the Spike region includes a segment of ~200 bases where the inferred divergence between RaTG13 and 2019-nCoV dramatically increases. This region is of interest as the spike protein is a determinant of viral host range and under heavy selection (14). The observed Spike region divergence indeed includes a substantial set of nonsynonymous differences (Supplemental Figure 1). Notably, while reads from the pangolin lung dataset mapped to this region do not show a similar increase in variation relative to the human 2019-nCoV, we also did not observe a significant drop in variance between human and pangolin in this region. While Wong et al. concluded that recombination likely occurred in the spike region in the derivation of 2019-nCoV, definitive conclusions regarding the origins of 2019-nCoV are difficult given the limited sequencing data available and without consideration to altered evolutionary rates in different lineages (15). Thus alternative models for the observed sequence variation seem most parsimonious, including that of selection acting on the RaTG13 sequences in bat or another intermediate host resulting in a rapid variation of this highly critical virus-receptor interface.

The availability of numerous paths (both targeted and agnostic) toward identification of natural niches for pathogenic sequences will remain useful to the scientific community and to public health, as will vigorous sharing of ideas, data, and discussion of potential origins and modes of spread for epidemic pathogens.

## Supporting information

SummaryOfMetasearchResults

## Authors’ contributions

All authors contributed equally, and author order was selected randomly.

---

**Supplemental Figure 1:**
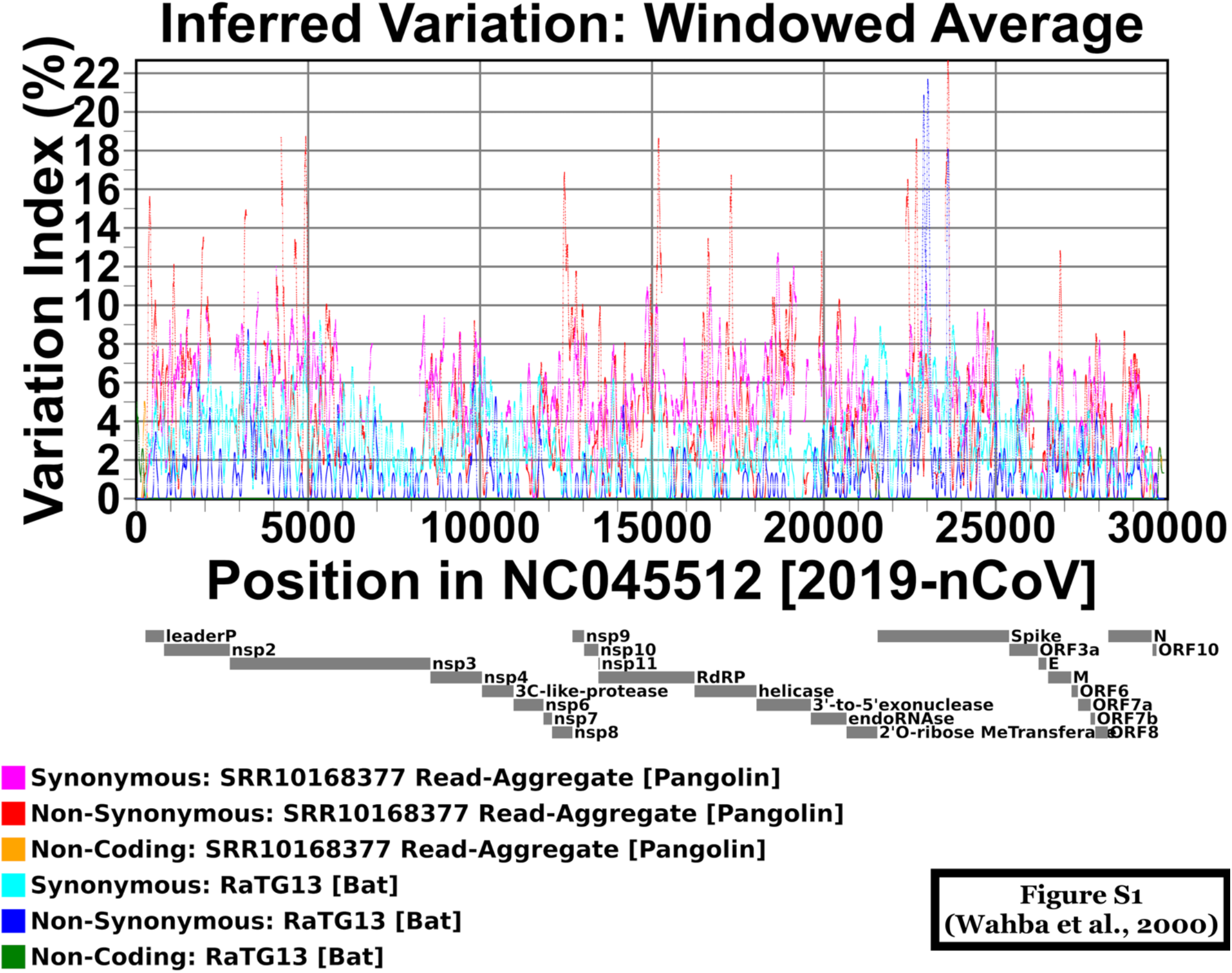
Detailed plot of inferred substitutions. The plot shows incidence of inferred synonymous, nonsynonymous, and noncoding substitution from comparison of RaTG13 assembly [2] and a pangolin coronavirus scaffold, to the 2019-nCoV isolate from Wu et al. [1]. The pangolin scaffold was generated from a Blast alignment of 2019-nCoV mapping reads in SRR10168377. This plot addresses several challenges associated with the limited sequencing data available by attempting to provide the most favorable alignment of that sequence possible. To maximize sensitivity in detecting potential recombination, ambiguities in which two or more reads apparently disagreed (which were rare; approximately 1.2% of assigned bases) were resolved in favor of “no substitution” at any position if one read matches the 2019-nCoV genome. This will provide a lower bound of variation, although regions covered by a single read are still subject to amplification and sequencing error. Near-perfect overlaps between reads from SRR10168377 argue that such error is relatively low as agreement in those regions is 99.6%. The cumulative count of synonymous, nonsynonymous, and noncoding variants are per base pair and are taken over a 75 bp window (approximately 1/2 the read length) and then averaged over each 75 bp window with a weighting inversely proportional to distance. This results in the observed inverted spike for each individual variant.

**TableS1.**
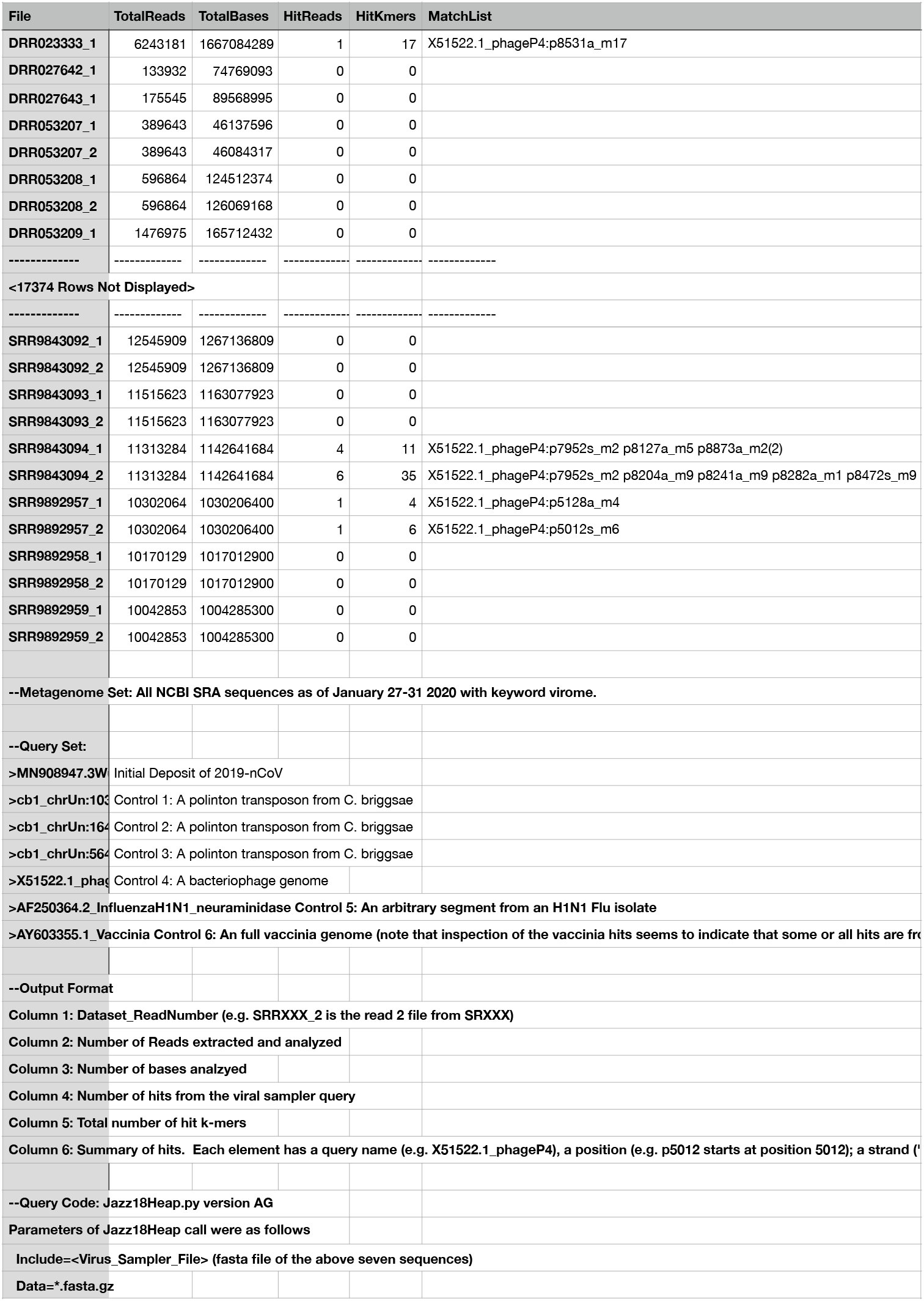
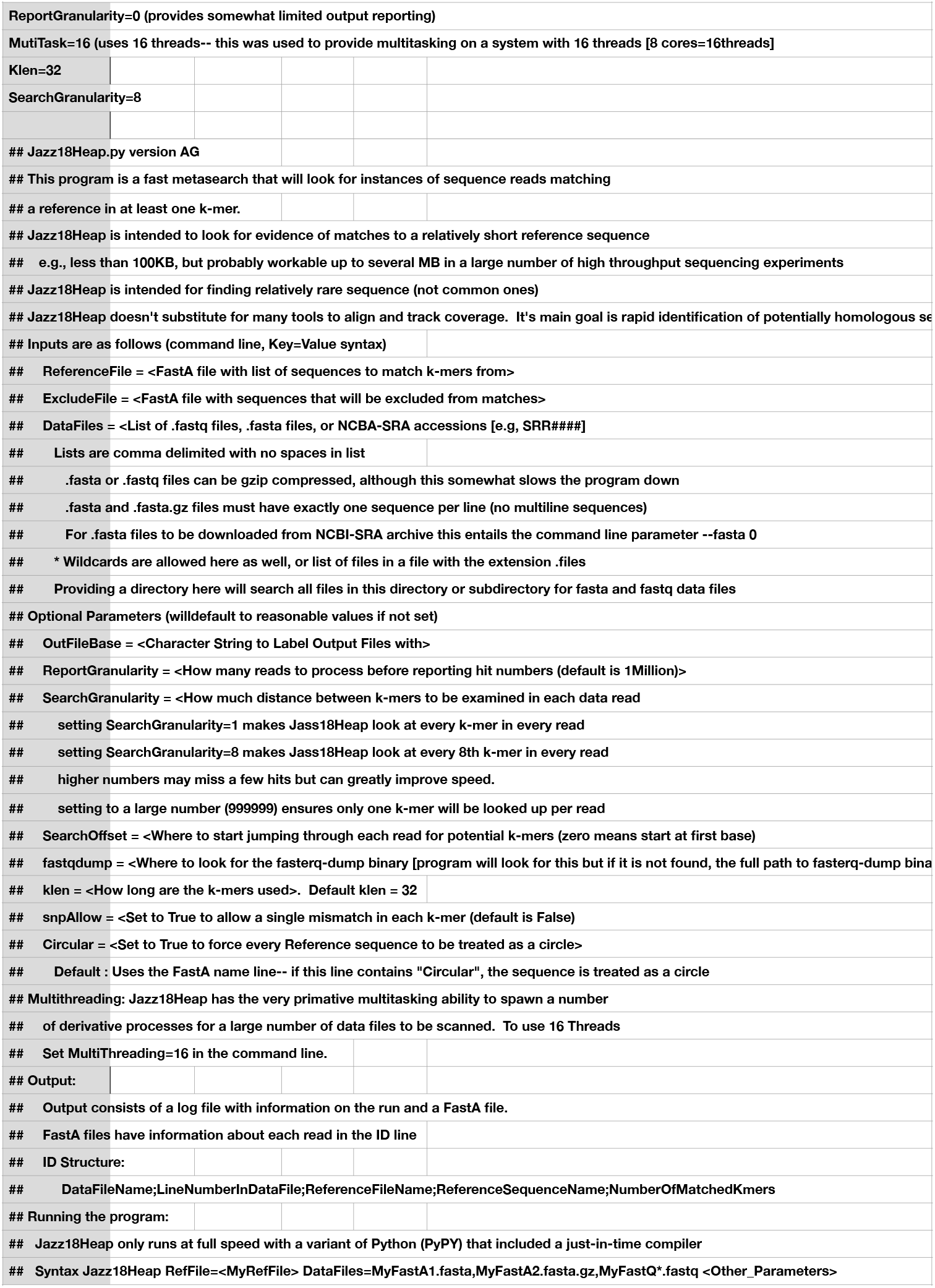
(Wahba et al. 2020) (Graphic below shows an abbreviated segment of Table S1, Full Table at GitHub.com/FireLabSoftware/MetaSearch)

| File | TotalReads | TotalBases | HitReads | HitKmers | MatchList |
| --- | --- | --- | --- | --- | --- |
| DRR023333_1 | 6243181 | 1667084289 | 1 | 17 | X51522.1_phageP4:p8531a_m17 |
| DRR027642_1 | 133932 | 74769093 | 0 | 0 |  |
| DRR027643_1 | 175545 | 89568995 | 0 | 0 |  |
| DRR053207_1 | 389643 | 46137596 | 0 | 0 |  |
| DRR053207_2 | 389643 | 46084317 | 0 | 0 |  |
| DRR053208_1 | 596864 | 124512374 | 0 | 0 |  |
| DRR053208_2 | 596864 | 126069168 | 0 | 0 |  |
| DRR053209_1 | 1476975 | 165712432 | 0 | 0 |  |
| --- | --- | --- | --- | --- | --- |
| <17374 Rows Not Displayed> |  |  |  |  |  |
| --- | --- | --- | --- | --- | --- |
| SRR9843092_1 | 12545909 | 1267136809 | 0 | 0 |  |
| SRR9843092_2 | 12545909 | 1267136809 | 0 | 0 |  |
| SRR9843093_1 | 11515623 | 1163077923 | 0 | 0 |  |
| SRR9843093_2 | 11515623 | 1163077923 | 0 | 0 |  |
| SRR9843094_1 | 11313284 | 1142641684 | 4 | 11 | X51522.1_phageP4:p7952s_m2 p8127a_m5 p8873a_m2(2) |
| SRR9843094_2 | 11313284 | 1142641684 | 6 | 35 | X51522.1_phageP4:p7952s_m2 p8204a_m9 p8241a_m9 p8282a_m1 p8472s_m9 |
| SRR9892957_1 | 10302064 | 1030206400 | 1 | 4 | X51522.1_phageP4:p5128a_m4 |
| SRR9892957_2 | 10302064 | 1030206400 | 1 | 6 | X51522.1_phageP4:p5012s_m6 |
| SRR9892958_1 | 10170129 | 1017012900 | 0 | 0 |  |
| SRR9892958_2 | 10170129 | 1017012900 | 0 | 0 |  |
| SRR9892959_1 | 10042853 | 1004285300 | 0 | 0 |  |
| SRR9892959_2 | 10042853 | 1004285300 | 0 | 0 |  |
| --Metagenome Set: All NCBI SRA sequences as of January 27-31 2020 with keyword virome. |  |  |  |  |  |
| --Query Set: |  |  |  |  |  |
| >MN908947.3W Initial Deposit of 2019-nCoV |  |  |  |  |  |
| >cb1_chrUn:103 Control 1: A polinton transposon from C. briggsae |  |  |  |  |  |
| >cb1_chrUn:164 Control 2: A polinton transposon from C. briggsae |  |  |  |  |  |
| >cb1_chrUn:564 Control 3: A polinton transposon from C. briggsae |  |  |  |  |  |
| >X51522.1_phageP4 Control 4: A bacteriophage genome |  |  |  |  |  |
| >AF250364.2 InfluenzaH1N1_neuraminidase Control 5: An arbitrary segment from an H1N1 Flu isolate |  |  |  |  |  |
| >AY603355.1 Vaccinia Control 6: An full vaccinia genome (note that inspection of the vaccinia hits seems to indicate that some or all hits are from the same genome) |  |  |  |  |  |
| --Output Format |  |  |  |  |  |
| Column 1: Dataset_ReadNumber (e.g. SRRXXX_2 is the read 2 file from SRRXXX) |  |  |  |  |  |
| Column 2: Number of Reads extracted and analyzed |  |  |  |  |  |
| Column 3: Number of bases analyzed |  |  |  |  |  |
| Column 4: Number of hits from the viral sampler query |  |  |  |  |  |
| Column 5: Total number of hit k-mers |  |  |  |  |  |
| Column 6: Summary of hits. Each element has a query name (e.g. X51522.1_phageP4), a position (e.g. p5012 starts at position 5012); a strand ('+' or '-') |  |  |  |  |  |
| --Query Code: Jazz18Heap.py version AG |  |  |  |  |  |
| Parameters of Jazz18Heap call were as follows |  |  |  |  |  |
| Include=<Virus_Sampler_File> (fasta file of the above seven sequences) |  |  |  |  |  |
| Data=*.fasta.gz |  |  |  |  |  |
| ReportGranularity=0 (provides somewhat limited output reporting) |  |  |  |  |  |
| MultiTask=16 (uses 16 threads-- this was used to provide multitasking on a system with 16 threads [8 cores=16threads]) |  |  |  |  |  |
| Klen=32 |  |  |  |  |  |
| SearchGranularity=8 |  |  |  |  |  |
| ## Jazz18Heap.py version AG |  |  |  |  |  |
| ## This program is a fast metasearch that will look for instances of sequence reads matching |  |  |  |  |  |
| ## a reference in at least one k-mer. |  |  |  |  |  |
| ## Jazz18Heap is intended to look for evidence of matches to a relatively short reference sequence |  |  |  |  |  |
| ## e.g., less than 100KB, but probably workable up to several MB in a large number of high throughput sequencing experiments |  |  |  |  |  |
| ## Jazz18Heap is intended for finding relatively rare sequence (not common ones) |  |  |  |  |  |
| ## Jazz18Heap doesn't substitute for many tools to align and track coverage. It's main goal is rapid identification of potentially homologous se |  |  |  |  |  |
| ## Inputs are as follows (command line, Key=Value syntax) |  |  |  |  |  |
| ## ReferenceFile = <FastA file with list of sequences to match k-mers from> |  |  |  |  |  |
| ## ExcludeFile = <FastA file with sequences that will be excluded from matches> |  |  |  |  |  |
| ## DataFiles = <List of .fastq files, .fasta files, or NCBI-SRA accessions [e.g, SRR###]> |  |  |  |  |  |
| ## Lists are comma delimited with no spaces in list |  |  |  |  |  |
| ## .fasta or .fastq files can be gzip compressed, although this somewhat slows the program down |  |  |  |  |  |
| ## .fasta and .fasta.gz files must have exactly one sequence per line (no multiline sequences) |  |  |  |  |  |
| ## For .fasta files to be downloaded from NCBI-SRA archive this entails the command line parameter --fasta 0 |  |  |  |  |  |
| ## * Wildcards are allowed here as well, or list of files in a file with the extension .files |  |  |  |  |  |
| ## Providing a directory here will search all files in this directory or subdirectory for fasta and fastq data files |  |  |  |  |  |
| ## Optional Parameters (will default to reasonable values if not set) |  |  |  |  |  |
| ## OutFileBase = <Character String to Label Output Files with> |  |  |  |  |  |
| ## ReportGranularity = <How many reads to process before reporting hit numbers (default is 1Million)> |  |  |  |  |  |
| ## SearchGranularity = <How much distance between k-mers to be examined in each data read |  |  |  |  |  |
| ## setting SearchGranularity=1 makes Jass18Heap look at every k-mer in every read |  |  |  |  |  |
| ## setting SearchGranularity=8 makes Jass18Heap look at every 8th k-mer in every read |  |  |  |  |  |
| ## higher numbers may miss a few hits but can greatly improve speed. |  |  |  |  |  |
| ## setting to a large number (999999) ensures only one k-mer will be looked up per read |  |  |  |  |  |
| ## SearchOffset = <Where to start jumping through each read for potential k-mers (zero means start at first base) |  |  |  |  |  |
| ## fastqdump = <Where to look for the fasterq-dump binary [program will look for this but if it is not found, the full path to fasterq-dump bina |  |  |  |  |  |
| ## klen = <How long are the k-mers used>. Default klen = 32 |  |  |  |  |  |
| ## snpAllow = <Set to True to allow a single mismatch in each k-mer (default is False) |  |  |  |  |  |
| ## Circular = <Set to True to force every Reference sequence to be treated as a circle> |  |  |  |  |  |
| ## Default : Uses the FastA name line-- if this line contains "Circular", the sequence is treated as a circle |  |  |  |  |  |
| ## Multithreading: Jazz18Heap has the very primitive multitasking ability to spawn a number |  |  |  |  |  |
| ## of derivative processes for a large number of data files to be scanned. To use 16 Threads |  |  |  |  |  |
| ## Set MultiThreading=16 in the command line. |  |  |  |  |  |
| ## Output: |  |  |  |  |  |
| ## Output consists of a log file with information on the run and a FastA file. |  |  |  |  |  |
| ## FastA files have information about each read in the ID line |  |  |  |  |  |
| ## ID Structure: |  |  |  |  |  |
| ## DataFileName;LineNumberInDataFile;ReferenceFileName;ReferenceSequenceName;NumberOfMatchedKmers |  |  |  |  |  |
| ## Running the program: |  |  |  |  |  |
| ## Jazz18Heap only runs at full speed with a variant of Python (PyPY) that included a just-in-time compiler |  |  |  |  |  |
| ## Syntax Jazz18Heap RefFile=<MyRefFile> DataFiles=MyFastA1.fasta,MyFastA2.fasta.gz,MyFastQ*.fastq <Other_Parameters> |  |  |  |  |  |

